# Human angiotensin-converting enzyme 2 transgenic mice infected with SARS-CoV-2 develop severe and fatal respiratory disease

**DOI:** 10.1101/2020.07.09.195230

**Authors:** Joseph W. Golden, Curtis R. Cline, Xiankun Zeng, Aura R. Garrison, Brian D. Carey, Eric M. Mucker, Lauren E. White, Joshua D. Shamblin, Rebecca L. Brocato, Jun Liu, April M. Babka, Hypaitia B. Rauch, Jeffrey M. Smith, Bradley S. Hollidge, Collin Fitzpatrick, Catherine V. Badger, Jay W. Hooper

**Affiliations:** Virology Division, United States Army Medical Research Institute of Infectious Diseases, Fort Detrick, MD 21702; Pathology, United States Army Medical Research Institute of Infectious Diseases, Fort Detrick, MD 21702; Diagnostic Systems Division, United States Army Medical Research Institute of Infectious Diseases, Fort Detrick, MD 21702; Comparative Medicine Division, United States Army Medical Research Institute of Infectious Diseases, Fort Detrick, MD 21702

## Abstract

The emergence of SARS-CoV-2 has created an international health crisis. Small animal models mirroring SARS-CoV-2 human disease are essential for medical countermeasure (MCM) development. Mice are refractory to SARS-CoV-2 infection due to low affinity binding to the murine angiotensin-converting enzyme 2 (ACE2) protein. Here we evaluated the pathogenesis of SARS-CoV-2 in male and female mice expressing the human ACE2 gene under the control of the keratin 18 promotor. In contrast to non-transgenic mice, intranasal exposure of K18-hACE2 animals to two different doses of SARS-CoV-2 resulted in acute disease including weight loss, lung injury, brain infection and lethality. Vasculitis was the most prominent finding in the lungs of infected mice. Transcriptomic analysis from lungs of infected animals revealed increases in transcripts involved in lung injury and inflammatory cytokines. In the lower dose challenge groups, there was a survival advantage in the female mice with 60% surviving infection whereas all male mice succumbed to disease. Male mice that succumbed to disease had higher levels of inflammatory transcripts compared to female mice. This is the first highly lethal murine infection model for SARS-CoV-2. The K18-hACE2 murine model will be valuable for the study of SARS-CoV-2 pathogenesis and the assessment of MCMs.

## Introduction

SARS-CoV-2 is a betacoronavirus and the causative agent of COVID-19, a febrile respiratory human disease that emerged in late 2019 in China and subsequently spread throughout the world (1, 2). COVID-19 is primarily a respiratory disease with a wide spectrum of severity ranging from a mild cough, to the development of hypoxia, and in some cases resulting in a life-threatening acute respiratory distress syndrome (ARDS) requiring mechanical ventilation (3, 4). The most severe cases are generally skewed towards the aged population (>50) and those with underlying health conditions such as hypertension or cardiovascular disorders (5, 6). SARS-CoV-2 human infections can also cause vasculature damage and coagulopathies, leading to infarction (7–9). Acute disease often presents with elevated levels of inflammatory cytokines, including IL-6, and these host molecules may play a role in the pathogenic process (10, 11). Additionally, ~30% of cases include signs of neurological disease such as headache, anosmia (loss of smell), ataxia, meningitis, seizures and impaired consciousness (12–14). To-date, SARS-CoV-2 has infected over eight million people world-wide and resulted in the death of more than 400,000. There is an urgent need for medical countermeasures to prevent this disease or limit disease severity in a post exposure setting.

Similar to SARS-CoV (15), SARS-CoV-2 binds to target cells via an interaction between the 139 kDa viral spike protein and the host angiotensin-converting enzyme 2 (ACE2) protein (16–19). An important factor in host tropism of the virus is this receptor interaction and reduced affinity between these two molecules greatly impacts host susceptibility to infection. Both SARS-CoV and possibly to a greater extent SARS-CoV-2 bind to murine ACE2 (mACE) poorly compared human ACE2 (hACE2) (20). Accordingly, mice are inherently refractory to infection by ACE2 utilizing human coronaviruses (21–23). In these animals, infection by SARS-CoV (22) and SARS-CoV-2 (23) is rapidly controlled, although older mice are more permissive to lung replication, but nevertheless lung injury is limited and mortality is generally low. Indeed, even infection in mice lacking adaptive immunity due to RAG deficiencies (no T-cells or B-cells) are protected against severe SARS-CoV disease, whereas mice genetically devoid of STAT-1, an important molecule involved in innate immunity, are sensitive to infection by SARS-CoV(24–26). However, disease in STAT-1 mice is protracted and not highly representative of human infections (25, 26).

In response to the need for small animal models to study SARS-CoV, several laboratories produced transgenic animals expressing the hACE2 gene under the control of various promotors, including the CMV, HFH4 epithelial cell promotor or the endogenous murine ACE2 promotor (27–31). Many of these animals develop severe disease upon infection by SARS-CoV, but severity largely depends upon expression levels and tissue distribution of the hACE2 transgene. One transgenic mouse strain, termed K18-hACE2, developed by Perlman S., et al. expresses the hACE2 molecule under the control of the keratin 18 promoter (30, 32). K18 limits expression to airway epithelial cells, colon and to a lesser extent kidney, liver, spleen and small intestine. A minor level of hACE2 expression was also detected in the brain. These mice are susceptible to SARS-CoV strain Urbani and develop severe respiratory disease subsequent to intranasal exposure characterized by lung inflammation, serum cytokine and chemokine production as well as high lethality (30). As with several mouse strains transgenic for hACE2, SARS-CoV infects the brains of K18-hACE2 mice (30, 32). CNS localization is speculated to play a major role in host mortality due to neuronal cell death, particular cell loss in cardiorespiratory control center (32). Some of these previous hACE2 transgenic mice are being used for SARS-CoV-2 research and newer systems have been developed using CRSIPR-Cas9 (23, 33, 34). However, none were shown to produce a consistent lethal disease and in models where lung injury occurred, it was most pronounced in aged animals (23, 34). Here, we evaluated susceptibility of K18-hACE2 mice to SARS-CoV-2. We found that mice developed severe disease that included respiratory distress with weight loss and mortality, as well as brain infection.

### SARS-CoV-2 produces severe and fatal infection of K18-hACE2 mice

Two groups of 9-week old K18-hACE2 mice (n=14 per group) were intranasally infected with 2×10^4^ or 2×10^3^ pfu of SARS-CoV-2. These groups equally divided by sex (n=7 per group/sex). On day 3, two mice per group were euthanized to assess disease severity. The remaining 5 mice per group were monitored up to 28 days. We also infected C57BL/6, BALB/c and RAG2 deficient mice with a challenge dose of 2×10^4^ pfu. On day 4, all groups of infected K18-hACE2 mice began to lose weight, which was more pronounced in the female mice compared to the male mice in either challenge dose group (**Fig. 1A & Fig. S1A**). Non-transgenic mice did not lose any weight and no animal succumbed to disease. Starting on day 5, several K18-hACE2 animals began to show signs of respiratory disease, included labored breathing and conjunctivitis. On day 5 through day 7 the majority of mice (15/20 mice) began to meet euthanasia criteria (n=13) or died (n=2). An additional animal in the low dose male group succumbed to disease on day 11, after a period of weight loss. The highest mean weight loss was in the female groups (>20%), although male mice lost >12% weight (**Fig S1A,B**). Of the K18-hACE2 mice in the high dose challenge groups, all females and 80% of the male mice succumbed to disease. Most female mice survived in the low dose group with 40% mortality, but all male mice succumbed to disease. The difference in survival between low dose male and female mice was significant (log-rank; p=0.040), as was mortality between the high and low dose female groups (log-rank; p=0.037). There was no statistical significance in survival between the other groups. Lung homogenates taken on day 3 showed high levels of virus in K18-hACE2, in contrast to C57BL/6 or RAG2-deficient mice which had low levels of virus (**Fig. 1B,C**). The virus RNA levels were nearly identical between all the infected K18-hACE2 groups and remained high in most of the euthanized mice, with the exception of the mouse that died on day 11. That animal had markedly lower levels of detectable genomic RNA (**Fig. 1C**). Compared to non-transgenic mice and uninfected K18-hACE2 mice, infected K18-hACE2 mice had comparatively higher serum levels of TNF-α, IL-6 and IL-10 as well as the monocyte chemoattractants MCP-1 (CCL2) and MCP-3 (CCL7) (**Fig. 1D**). Levels of cytokines observed in mice that succumbed to disease were generally higher compared to those sampled on day 3. Overall, these findings indicated that SARS-CoV-2 causes a severe disease in K18-hACE2 mice following intranasal exposure.

**Figure 1.**
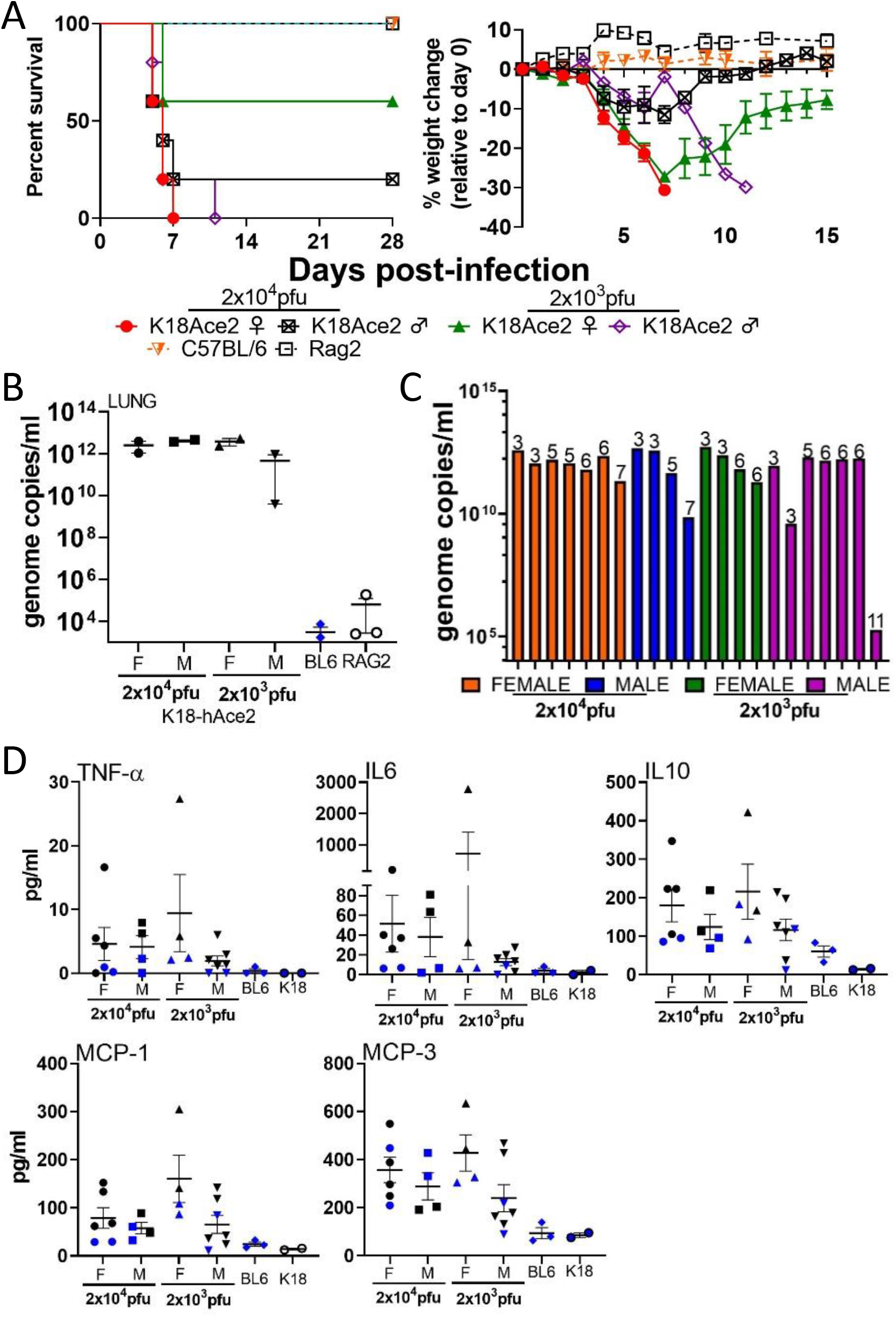
SARS-CoV-2 infection in K18-hACE2 transgenic mice. **A.** Male and Female K18-hACE2 transgenic mice (Day 0-3 n=7/group; Day 3+, n=5/group) were infected with 2×10^4^ PFU or 2 × 10^3^ PFU of SARS-CoV-2 by the IN route. C57BL/6 and RAG2 KO mice (Day 0-3 n=8/ group; Day 3+, n=5/group) were infected with 2×10^4^ pfu by the IN route. Survival and weight loss (+/− SEM) were monitored and plotted using Prism software. **B**. Titers in lung (n=2 mice/group) were examined on day 3 by qRT-PCR. Mean titers +/− SEM of the genome molecules of viral RNA/ml were graphed. **C.** Titers in lungs of individual K18-hACE2 mice. Numbers above bars denote day of death. Colors represent the four groups. **D.** Monocyte chemoattractants and inflammatory cytokines were measured from the serum of SARS-CoV-3 infected mice on day 3 or at the time of euthanasia using a multiplex system. Mice from each group are aggregated from samples taken on day 3 (blue symbols) and when mice were euthanized (black symbols).

### Lungs of K18-hACE2 mice exposed to SARS-CoV-2 show signs of acute disease

Lungs were collected from K18-hACE2 on day 3 or at the time of euthanasia due to disease severity. In K18-hACE2, viral RNA was detected by in situ hybridization (ISH) in all mice on day 3 and in most mice succumbing to disease on days 5-7, but these levels diminished as disease progressed (**Fig 2A, Fig S2, Table S1**). Additionally, ISH labeling was patchy (**Fig. S2**) and most severe at the day 3 time point. ISH labeling was present in both inflamed and normal appearing alveolar septa. Positive ISH labeling for SARS-CoV-2 was identified multifocally in alveolar septa in the lung, suggesting infection of pneumocytes and macrophages. This was confirmed by the detection of SARS-CoV-2 protein in E-cadherin positive cells (pneumocytes) and CD68 positive (alveolar and infiltrating macrophages) using immunofluorescence assay (IFA, **Fig. 2B,C**).

**Figure 2.**
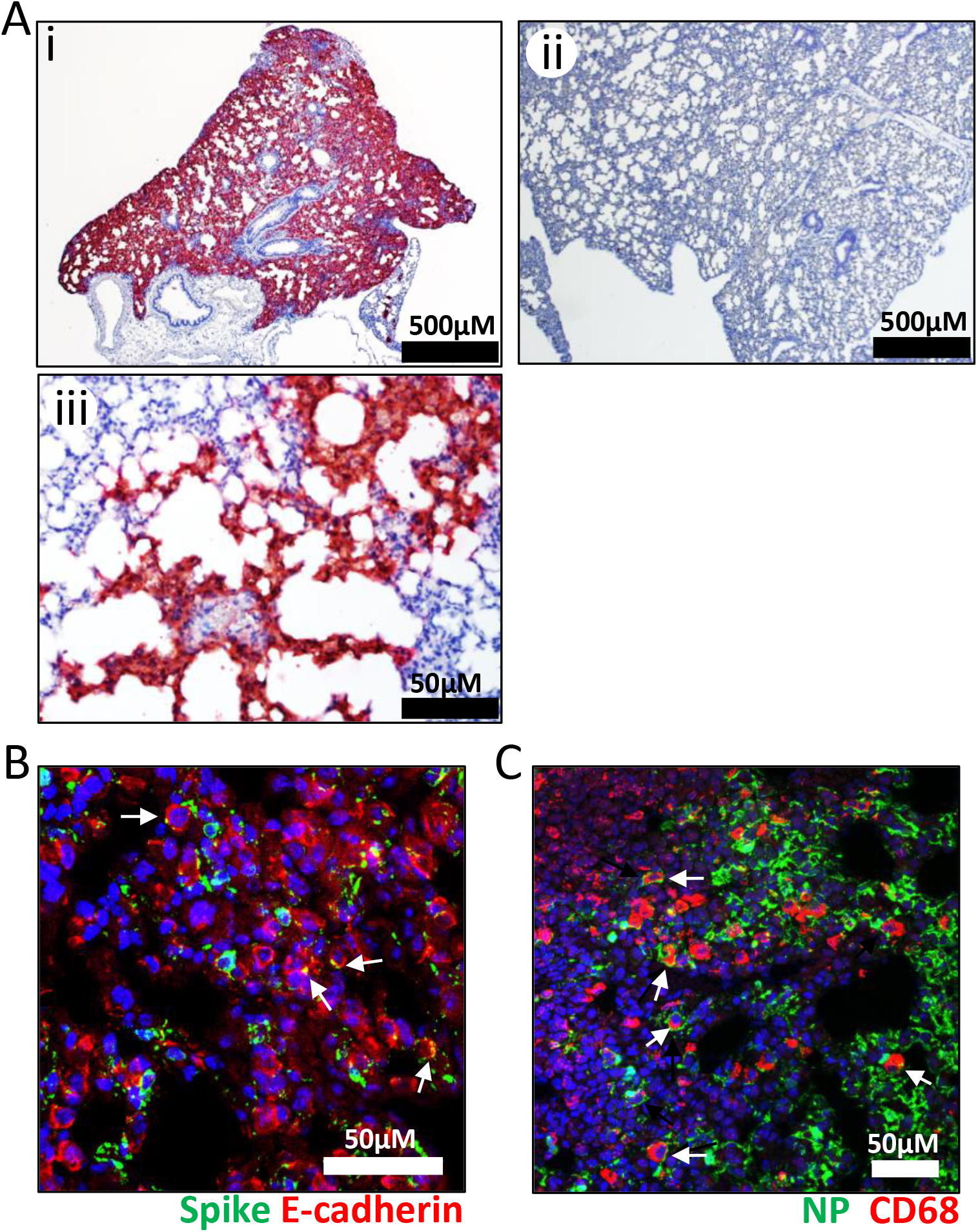
Infection of SARS-CoV-2 in the lung of K18-ACE2 transgenic mice. **A.** Representative ISH images showing the presence of SARS-CoV-2 RNA (red) in the lungs of infected at low and high magnification (i & iii) or uninfected (ii) K18-hACE2 mice. Cells were counterstained with hematoxylin (blue). **B.** Co-staining for viral spike protein (green) and E-cadherin (red) in infected lung tissues using IFA. Arrows point to double positive cells. Nuclei are stained with DAPI (blue). **C.** Co-staining of viral nucleoprotein and the macrophage marker CD68 (red) in infected lungs using IFA. Arrows denote double positive cells. Nuclei are stained with DAPI (blue).

In comparison with the normal lung architecture in uninfected control animals, infected mice necropsied on day 3, and those succumbing to disease on days 5-11, had varying levels of lung injury including area of lung consolidation characterized by inflammation/expansion of alveolar septa with fibrin, edema and mononuclear leukocytes and infiltration of vessel walls by numerous mononuclear leukocytes (**Fig 3A, Fig S4, and Table S1**). Type II pneumocyte hyperplasia was identified in less than half of infected animals. This lesion had a relatively patchy distribution except in the most severely affected animals where it is more abundant. In areas of septal inflammation, exudation of fibrin and edema into alveolar lumina from damaged septal capillaries was observed in most animals. Vasculitis was the most common finding and was present in ~95% percent of all mice (**Fig. 3A,B**). The lesion encompassed small and intermediate caliber vessels, and was often characterized by near circumferential infiltration of the vessel wall by numerous mononuclear inflammatory cells. This lesion also contained small amounts of fibrin and occasional necrotic debris, affecting all tunics and obscuring the vessel wall architecture. However, the endothelial cells surrounding lung vasculature were largely free of viral RNA (**Fig. 3B**). In one low dose male mouse that died on day 11, evidence of numerous fibrin thrombi were identified in the small and intermediate vessels, suggestive of a hypercoagulable state within the lung. This same animal had marginally detectable virus in the lung (**Fig. 1C**), and fibrin thrombi were not identified in other organs including liver and kidney. Infected K18-hACE2 mice had elevated numbers of TUNEL positive cells, suggesting increased cell death (**Fig. 3C**). We also detected increased expression of Ki-67, a marker for cellular proliferation, likely expressed by proliferating pneumocytes and potentially by replicating alveolar macrophages (**Fig. 3D**). Neutrophilia was detected by H&E staining, consistent with an increase in myeloperoxidase (MPO)-positive cells (neutrophils, basophil and eosinophil) detected by IFA (**Fig. S3A**). However, the MPO positive cells were devoid of viral antigen (**Fig. S3A**). There was also an increase in the presence of CD68 positive macrophages (**Fig. S3B**) and a pronounced increase in CD45 and CD3 positive cells in infected lungs, indicative of infiltrating leukocytes including T-cells, which is consistent with histologic findings (**Fig. S3C**). These data indicate that K18-hACE2 mice develop a pronounced lung injury upon exposure to SARS-CoV-2.

**Figure 3.**
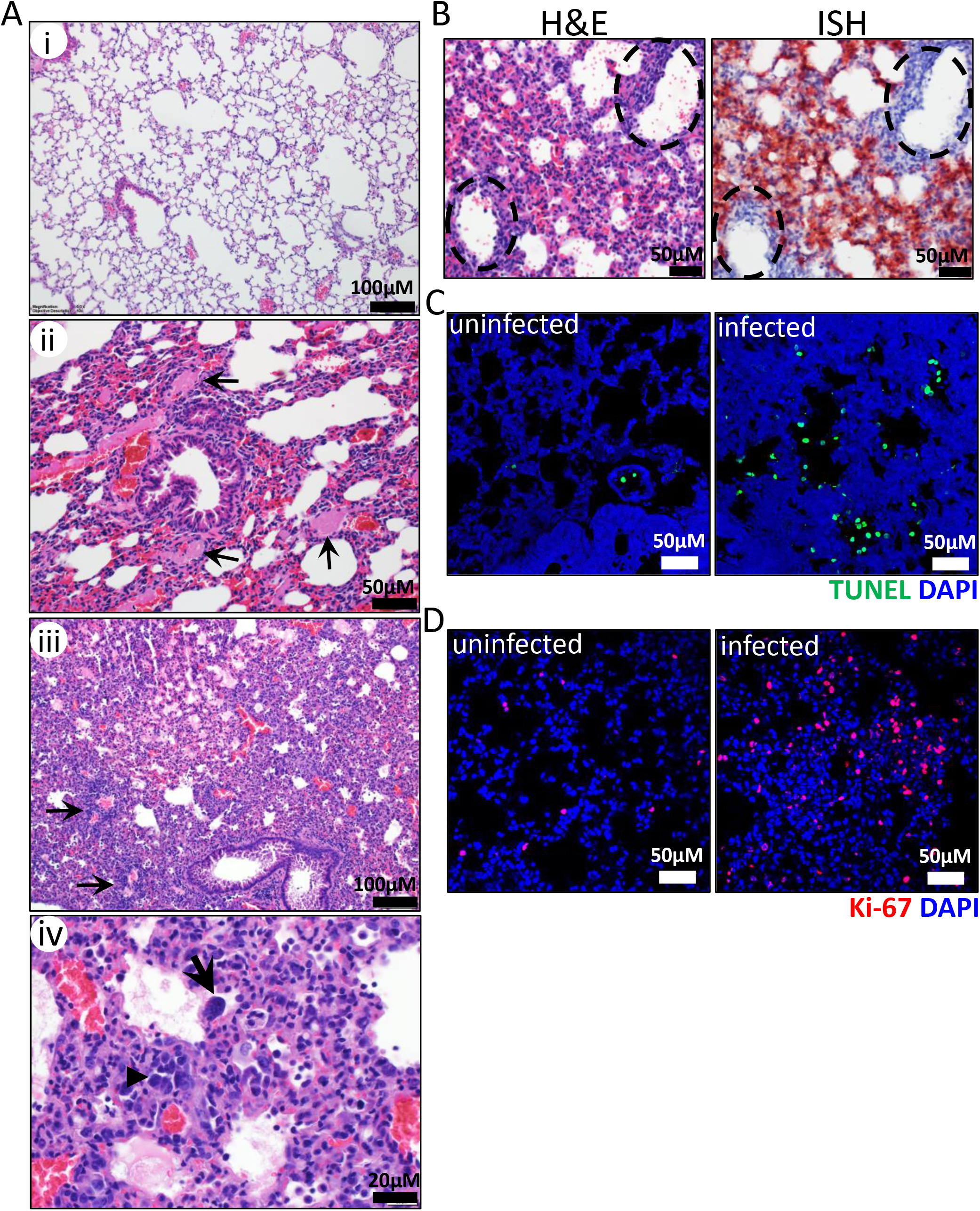
SARS-CoV-2 infection causes respiratory damage in K18-hACE2 mice. **A.** Representative H&E staining of lungs in infected C57BL/6 mice (i) or K18-hACE2 infected mice (ii-iv). Numerous fibrin thrombi (black arrows) filling the lumen of small to intermediate sized vessels (ii) adjacent to a normal bronchus and surrounded by minimally inflamed, congested and collapsed alveolar septa. Extensive area of lung consolidation (iii) with inflammation/expansion of alveolar septa, exudation of fibrin and edema into alveolar lumina and infiltration of vessel walls and perivascular area by numerous mononuclear inflammatory cells (arrows), disrupting/obscuring vessel architecture (vasculitis). Extensive area of consolidated lung (iv) showing type II pneumocyte hyperplasia (arrowhead) and rare multinucleate cells (black arrow), **B**. H&E and ISH staining of infected mouse lung showing vasculitis with absence of viral RNA in the affected vessel walls; note there is viral RNA (red) in the adjacent alveolar septa. Vessels are highlighted by the broken black circles. **C**. TUNEL staining of infected and uninfected K18-hACE2 mouse lungs. TUNEL (green) was performed as indicted in the materials and methods. Cell nuclei are stained with DAPI (blue). **D**. Ki-67 staining (red) in infected and uninfected K18-hACE2 mouse lungs. Nuclei are stained with DAPI (blue).

Transcriptional profiles of host immunological and inflammatory genes in lung homogenates from SARS-CoV-2 infected K18-hACE2 and C57BL/6 mice were examined by NanoString-based gene barcoding on day 3 (K18-hACE2 and C57BL/6) or in K18-hACE2 mice at the time euthanasia. 478 transcripts were increased in K18-hACE2 mice and 430 decreased at a log2 fold cutoff of 1 and a p value <0.05. Many of the increased transcripts in the K18-hACE2 mice were inflammatory genes including IL-6, interferon gamma and chemokines (CCL2, CCL5, CCL9 and CXCL10) (**Fig. 4A**). The type I interferon transcripts IFNA1 and IFNB1 along with the cytokines IL-9 and IL-2 were decreased. Consistent with the increased presence of CD68 macrophages, CD68 transcripts were also significantly increased in K18-hACE2 mice. Viral sensing pathways were elevated in infected mice with severe disease, indicated by high transcript levels of Irf1, RIG-I (Ddx58) and MDA-5 (Ifih1). Among the highest activated transcripts in the K18-hACE2 mice were genes involved in lung injury and hypoxia, including Sphingosine-1-Phosphate Lyase 1 (Sgpl1), Resistin-like molecule alpha (Retnla, FIZZ and HIMF), stromal cell-derived factor 1 (SDF1/CXCL12) and Hypoxia-inducible factor 1-alpha (HIF1A). No major difference between the challenge doses were observed. There were 13 genes increased and 21 decreased between the male and female mice (**Fig. 4B**). Male mice had higher levels of inflammatory cytokine transcripts including CXCL2, IL6, IL1R2 and CXCL1. Transcripts expressed more highly in female animals included Fatty acid synthase (Fasn) and Histone demethylase UTY (Uty). Collectively, these findings indicate that K18-hACE2 mice develop transcriptomic signatures of lung injury after exposure to SARs-CoV-2 with increased expression of inflammatory genes, inflammatory gene transcripts and markers of lung injury and hypoxia.

**Figure 4.**
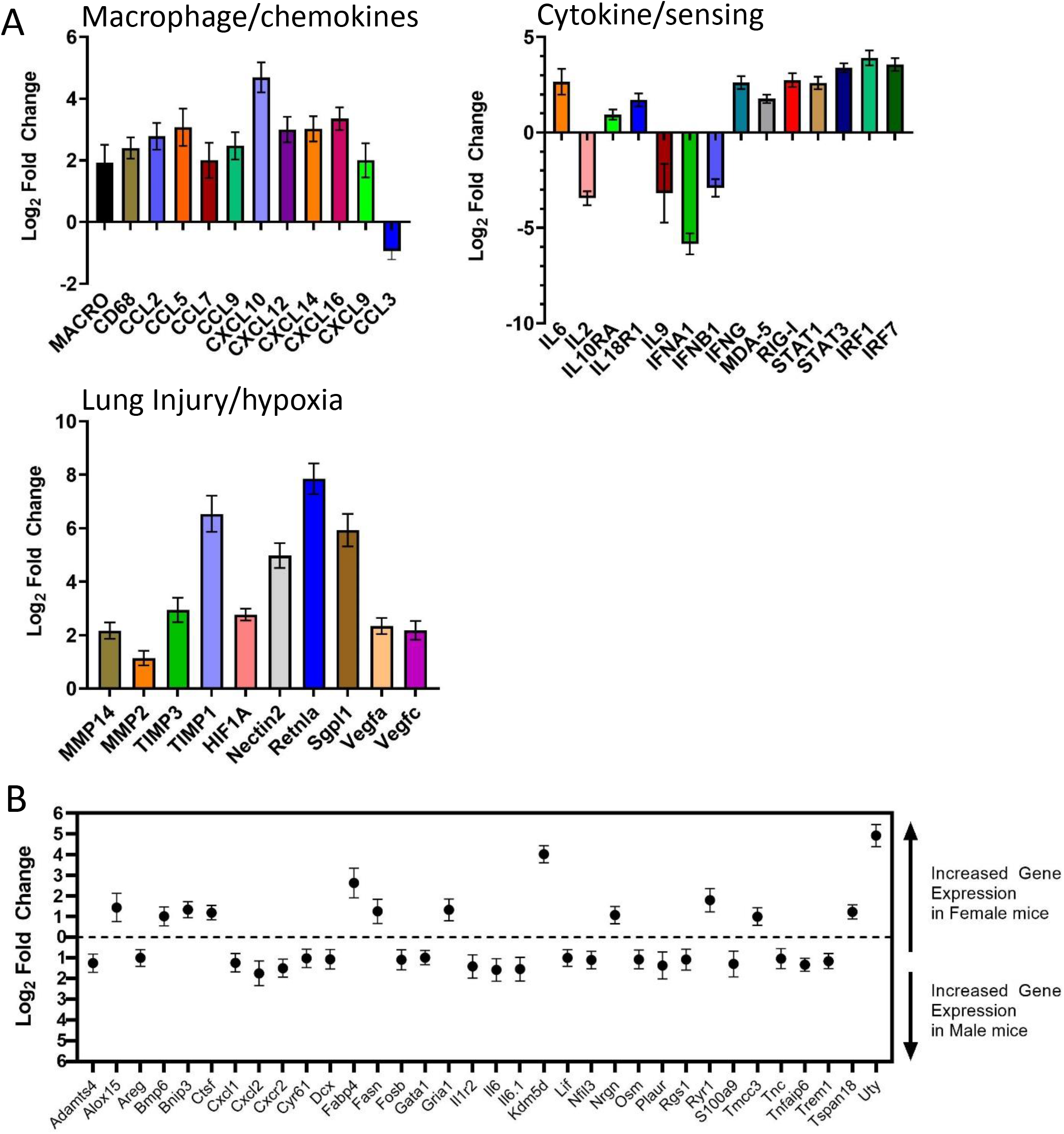
Transcriptional activation in SARS-CoV-2 infected lungs. Transcriptional activation in lung homogenates from infected K18-hACE2 (male and female) and C57BL/6 (female) mice were examined by NanoString. **A.** Log_2_ fold changes in gene expression levels of selected genes categorized by group versus infected C57BL/6 mice were graphed with SD. All graphed transcripts had a p value of <0.05. **B.** Differential gene expression (Log_2_ changes) between infected male and female K18-hACE2 mice. All graphed transcripts had a p value of <0.05.

### SARS-CoV-2 is present in the nasal cavity and eyes of K18-hACE2 mice

SARS-CoV-2 RNA was observed in the nasal turbinates in the majority of mice and rarely within the eyes of infected mice (**Fig. 5A, Fig. S5 and Table S1**). Viral RNA in the eye was localized to the retina, suggesting viral infection of neurons in the inner nuclear layer and ganglion cell layer (**Fig. S5**). Despite infection, evidence of inflammation or other damage in the retina or elsewhere in the eye was not present. Viral RNA and spike protein were also detected to some degree in the nasal turbinate epithelium (predominantly olfactory epithelium) as early as day 3 post-infection (**Fig. 5B**), as indicated by co-staining with cytokeratin (**Fig 5C**). Pathology was minimal, predominantly isolated to few areas of olfactory epithelium atrophy, with degeneration or erosion present in the epithelium lining the dorsal and lateral nasal meatuses (**Fig. 5D**). Some cellular sloughing was also detected and these sloughed cells contained viral RNA (**Fig. S5**). In mice succumbing to disease on days 5-7, virus was present in the olfactory bulb and most animals showed asymmetrical staining, with one bulb more positive than the other. Viral RNA was detected throughout the olfactory bulb, including in the olfactory nerve layer (ONL), glomerular layer (GL), external plexiform layer (EPL), and mitral cell layer (MCL) of olfactory bulbs in most of animals. Viral protein co-localized with the neuronal marker NeuN, suggesting virus was present within neurons in the olfactory bulb (**Fig. S6**). Virus was not detected in the olfactory bulb of animals taken on day 3, suggesting that virus trafficked to this region on day 4 or 5. These data indicated that SARS-CoV-2 infects cells within the nasal turbinates, eyes and olfactory system and that infection was observed in epithelial cells and neurons.

**Figure 5.**
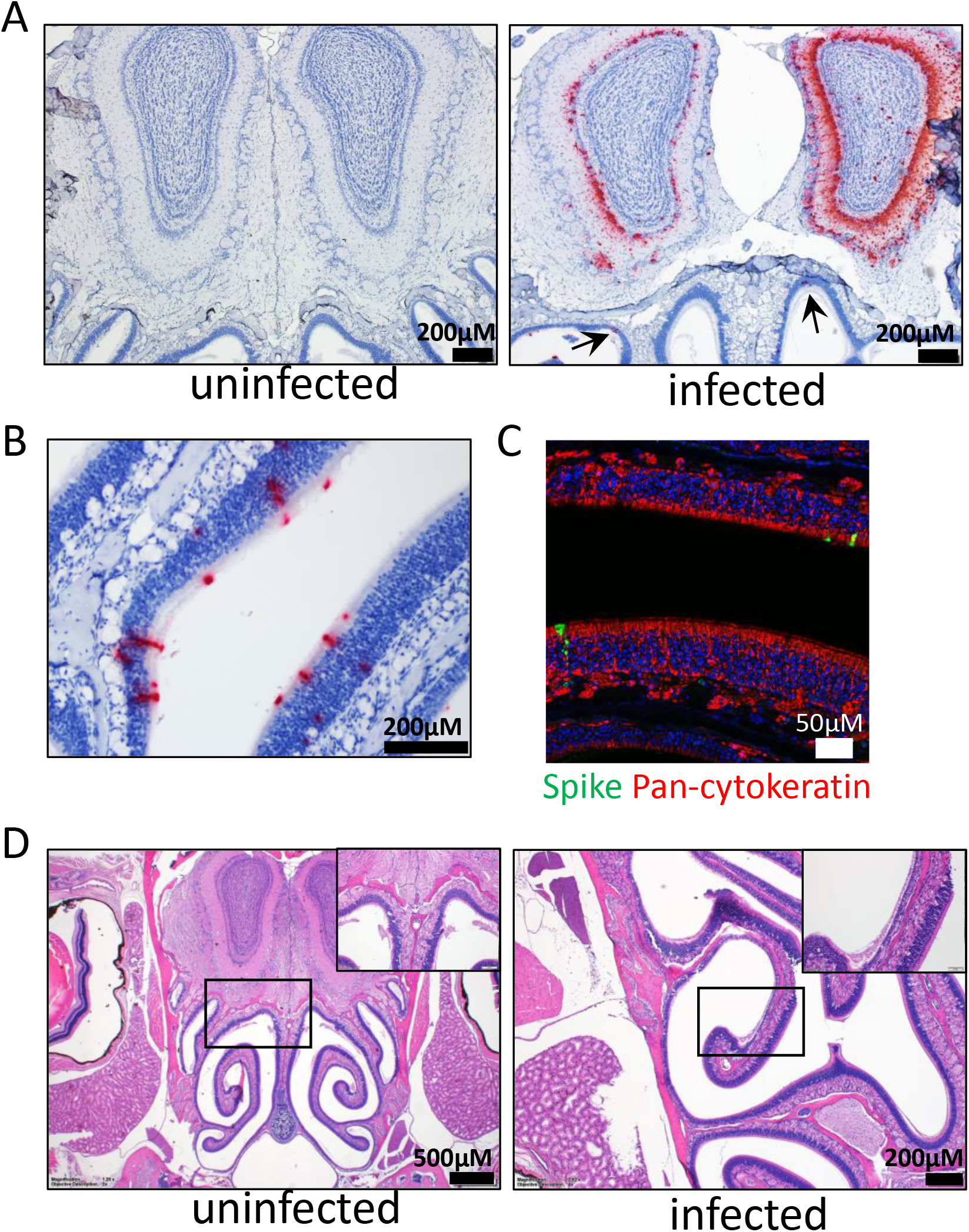
Infection of the nasal cavity and olfactory bulb. **A.** ISH labeling for SARS-CoV-2 RNA in a coronal section of the head including the caudal aspect of the nasal cavity and olfactory bulb. Within the olfactory bulb, a strong positive signal is present in the glomerular layer, external plexiform layer, mitral cell layer and internal plexiform layer, with multifocal positivity in the granular cell layer in the olfactory bulb hemisphere at right. Low numbers of cells within the olfactory epithelium lining the dorsal nasal meatus have a positive ISH signal (arrows). Cells were counterstained with hematoxylin (blue). **B**. ISH labeling for SARS-CoV-2 RNA in the olfactory epithelium. Cells were counterstained with hematoxylin (blue). **C**. IFA of olfactory epithelium stained for SARS-CoV-2 spike (green) and Pan-cytokeratin (red). Nuclei were stained with DAPI (blue). **D.** Representative H&E staining of the nasal cavity including olfactory epithelium and olfactory bulb from uninfected or infected K18-hACE2 mice. In the infected mouse there is a focal area of olfactory epithelium atrophy on a nasal turbinate located in the lateral nasal meatus. Inset image shows the indicated area of detail under increased magnification.

### SARS-CoV-2 infects the brain of K18-hACE2 mice

Brain infection was not observed in the majority of animals examined on day 3, but was prevalent in mice necropsied on days 5-11 (**Table S1**). Evidence of SARS-CoV-2 was found throughout the brain including strong but variably diffuse signal in regions of the thalamus, hypothalamus, amygdala, cerebral cortex, medulla, pons and midbrain (**Fig. 6A, Table S1**). Similar intense but less diffuse signal was present within the hippocampus. ISH positive cells included neurons of thalamic nuclei. In contrast, cells within the vessel walls and perivascular spaces infiltrated by mononuclear inflammatory cells were negative for viral genomic RNA (**Fig. 6A**). Histopathological changes were detected in the brains of several infected K18-hACE2 mice euthanized on day 5-11, but not in most mice taken on day 3 (**Fig. 6B, Table S1 and Fig. S7**). In the thalamus/hypothalamus, vasculitis was the most common lesion characterized by endothelial hypertrophy and increases in mononuclear leukocytes within the vessel wall and/or filling the perivascular space. Small amounts of necrotic debris were also identified. In the majority of these cases, the vascular lesion was characterized by the presence of increased numbers of microglia within the adjacent neuropil. Occlusive fibrin thrombi were also detected within the thalamus in a few mice. Meningitis was observed in a subset of animals and was associated with infiltration of mononuclear leukocytes (majority lymphocytes) and is most prominent adjacent to vessels. In the mouse that died late on day 11 with massive pulmonary clotting, the rostral cerebral cortex brain lesions included small to intermediate size vessel walls multifocally expanded by mononuclear inflammatory cells and perivascular hemorrhage extending into the adjacent neuropil (**Fig. 6B**). Increased numbers of microglia were readily detected on H&E stained sections, expanding outward from the perivascular neuropil. An increase in numbers of microglia were found in the neuropil surrounding affected vessels. Additionally, brains also showed signs of neuroinflammation indicated by increased staining of Iba-1 and GFAP indicating microgliosis and astrocytosis, respectively (**Fig. 6C**). Necrosis was identified in at least five animals, and was most prominent within the periventricular region of the hypothalamus as well as in the amygdala. The lesion was characterized by moderate numbers of shrunken, angular cells with hypereosinophilic cytoplasm, pyknotic nuclei and surrounded by a clear halo **(Fig. S7**). The morphology and location of individual cells was suggestive of neuronal necrosis, but further investigation will be required to confirm this finding. Viral spike protein was detected in NeuN positive cells, indicating viral infection of neurons (**Fig. 6D**). Viral antigen was absent in GFAP positive cells suggesting virus does not productively infect astrocytes. These data indicate that similar to SARS-CoV, SARS-CoV-2 also targets the brain of K18-hACE2 mice, causing brain injury. As indicated by duplex ISH labeling of brain, neurons are positive for both hACE2 transgene expression and viral genomic RNA (**Fig. S8**). No animal showed outward signs of neurological deficient, such as hind limb paralysis, head tilting or tremors.

**Figure 6.**
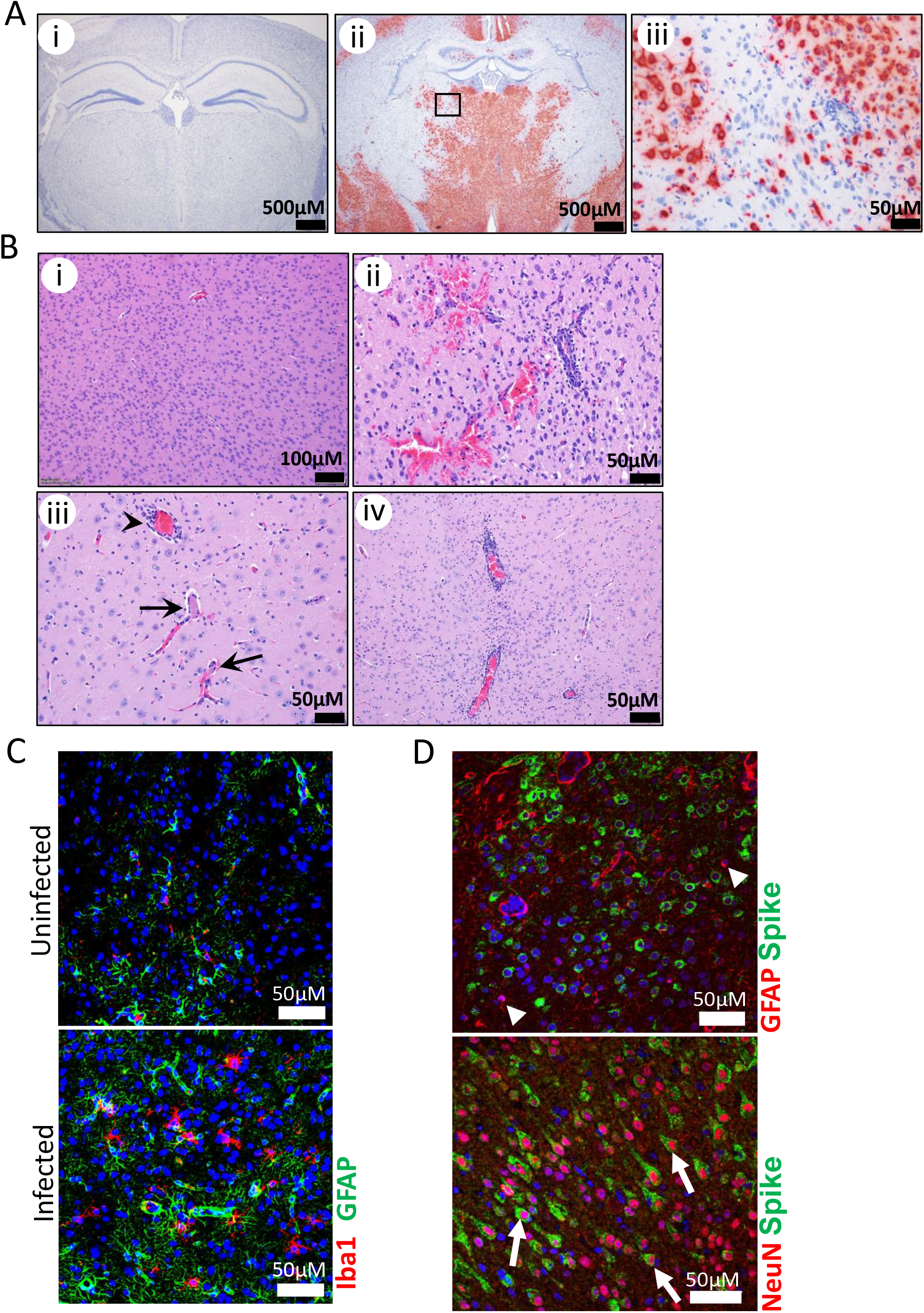
Neuropathogenesis of SARS-CoV-2 in K18-ACE2 transgenic and non-transgenic mice. **A.** ISH detection of for SARS-CoV-2 RNA in uninfected (i) and infected mice (ii and iii) in a coronal section of brain demonstrating a strong positive signal within neurons of thalamic nuclei. The boxed (ii) area is shown at increased magnification in the right panel (iii). Note the absence of a positive signal from the vessel at center right (iii) where the vessel wall and perivascular space are infiltrated by mononuclear inflammatory cells. **B.** Representative H&E staining of uninfected (i) or infected (ii-iv) K18-hACE2 mice. Perivascular hemorrhage extending into the adjacent neuropil (ii), in the region of the rostral cerebral cortex. Two small caliber vessels (iii) in the thalamus with fibrin thrombi (arrows), with mild microgliosis and some perivascular hemorrhage (arrowhead). The walls of small to intermediate size vessels and perivascular spaces (iv) are multifocally expanded/obscured by mononuclear inflammatory cells and increased numbers of glial cells **C.** Detection of Iba-1 and GFAP, markers for microgliosis and astrogliosis, respectively, in uninfected or infected brain sections using IFA. Nuclei were stained with DAPI (blue). **D.** Co-staining for astrocyte marker GFAP (red) or neuron marker NeuN (red) with SARS-CoV-2 spike protein (green). Spike protein was predominantly detected in NeuN^+^ neurons. Nuclei were stained with DAPI (blue).

## DISCUSSION

Other murine infection models for SARS-CoV-2 involving transgene expression of the human ACE2 protein have been reported (23, 33, 34). However in these models, SARS-CoV-2 only produces a transient weight loss with some lung injury, but the animals generally recover. Additionally, several of these models required mice aged >30 weeks for the most severe disease to occur, diminishing the practicality of these systems given the urgent need for medical countermeasures (MCMs) (23, 34). One system tested SARS-CoV-2 infection in mice in which hACE2 was expressed under the control of the HFH4 promoter (33). Infection in these mice was only ~40% lethal (2/5 mice) and lung injury (assessed by plethysmography) and weight loss were absent. Lethality in this system was exclusive to animals where virus was detected in the brain. Other recently reported SARS-CoV-2 murine models involved transduction of mouse lungs with a replication-incompetent Adenovirus virus or an adeno associated virus (AAV) encoding the hACE2 gene (35, 36). Transduced lung cells expressing hACE2 supported SARS-CoV-2 replication and lung pathology ensued along with weight loss. However, disease was generally mild with no lethality. Blockade of the type I interferon system using pharmacological intervention was needed to produce the most severe disease^26^. Murine systems faithfully producing the major elements of severe disease observed in humans will be more useful for identifying the most promising MCMs. The K18-hACE2 mice lost considerable weight >12% in males and >20% in females and lethality in the high dose exceeded 90%. Acute lung injury was detected in all animals succumbing to disease, with vascular damage the most common lesion. Similar to findings with SARS-CoV, SARS-CoV-2 infected the brains of K18-hACE2 mice. Brain infection resulted in vasculitis and inflammation, with SARS-CoV-2 antigen detected in neurons. It is possible virus enters the brain via the olfactory bulb, as has been reported for SARS-CoV, although more studies will be required as virus may also have entered the brain via inflamed vessels. Whether mortality results directly from brain infection is not clear, but this has been speculated as the major cause of mortality in SARS-CoV infected mice (32). Infection in the brain was delayed by at least four days, as it was an uncommon finding in day 3 animals. Thus, early during infection lung appears to be the primary target. We have not yet evaluated the protective efficacy of MCMs in this model, but it has been reported that antibodies protect against SARS-CoV, demonstrating the K18-hACE2 system is useful for evaluating countermeasures against ACE2-targeting human coronaviruses. Importantly, in our study some animals at the lower dose survived infection despite significant weight loss (>20% in female mice), indicating the model is not hypersensitive to infection and animals can recover. The K18-hACE2 animal model is commercially available and thus should provide an important platform to evaluated MCMs against SARS-CoV-2.

Infection of K18-hACE2 mice by SARS-CoV-2 produces a disease similar to that observed in acute human cases, with development of an acute lung injury associated with edema, production of inflammatory cytokines and the accumulation of mononuclear cells in the lung. Impacted lungs had elevated levels of transcripts consistent with respiratory damage such as increased expression of HIMF, which is involved in activation of lung endothelial cells in response to lung inflammation (37). We also found increased levels of Sgpl1, a molecule associated with lung injury (38) and known to be increased in mechanically ventilated mice (39). A prominent finding in infected K18-hACE2 mice system was vasculitis. Endotheliitis/vasculitis has been observed in human COVID-19 patients and a role for the endothelium in acute disease is becoming more apparent (7, 8, 40). Direct viral infection in human endothelial cells has also been reported (7). Curiously in the mice, virus was absent in these areas suggesting vasculitis was due to host inflammatory processes, but further study will be required to determine if vascular damage is incurred by direct or indirect viral effects. It has been speculated that during human acute disease, inflammatory cytokines (IL-6 in particular) are major drivers of tissue damage and this is supported by data from humans showing IL-6 receptor targeting with pharmacological antagonists (tocilizumab) can reduce morbidity (41). We found IL-6 transcripts, as well chemokines, are elevated in mouse lungs. During human disease, pulmonary inflammation is associated with increases in lung granulocytes and an increase in macrophages (4, 42, 43). Some have speculated that these macrophages play an important role in host injury (44). It is still unclear if human macrophages are directly infected by SARS-CoV-2, though we found virus antigen in CD68 macrophages and others report infection in murine MAC2-positive macrophages (23). Further study will be required to determine the role macrophages play in SARS-CoV-2 lung injury and given the regents available, murine systems may be highly suited for these studies. In addition to vascular issues, coagulopathies are a common findings in humans (9), with pulmonary embolism having been reported, along with clotting abnormities leading to loss of limbs (45). At least some mice produced evidence of these clotting issues, with one mouse presenting with fibrin thrombi in the lungs.

During human infections, males have been reported to have more severe outcomes despite a similar infection rate (46). In our experiments, we observed a statistically significant difference in survival of female and male mice infected at the lower dose of virus, with 60% of females but no males surviving infection. Despite surviving, the female mice challenged with the low dose still lost a significant amount of weight (>20%). Transcriptomic profiling in mouse lungs indicated that female mice that succumbed to disease had modestly lower levels of IL-6, CXCl-2 and IL-1RA suggesting a less intense inflammatory response. Our work only involved a small number of animals, and more work will be required to determine if the K18-hACE2 system recapitulates this sex difference in disease severity.

The neuroinvasive aspects of COVID-19 are becoming more appreciated(12). Indeed, SARS-CoV-2 causes neurological sequela in at least a third of human cases including headache, confusion, loss of smell, meningitis, seizures and ischemic and hemorrhagic stroke. The pathophysiology of human neurological injury induced by SARS-CoV-2 is ambiguous and a direct role of the virus has not been established. Other coronaviruses are associated with neurological disease, including HEV67N in porcine, and in humans SARS-CoV and MERS-CoV(47). SARS-CoV-2 infected K18-hACE2 mice showed evidence of neuroinflammation, neurovasculitis, meningitis and hemorrhage, emulating many of the human findings. SARS-CoV has been shown to infect the human and mouse brains and data suggest it replicates in neurons. In the K18-hACE mice, neuronal cells in the olfactory bulb, retina and brain were positive for SARS-CoV-2 viral RNA and antigen, suggesting these cells are supportive of a productive infection. In the K18-hACE2, we suspect virus accesses the brain via infection of the olfactory bulb and then spreads to neurons and other regions of the brain via connective neuron axons (32). In humans virus may gain access to the brain by infection of peripheral neurons, such as those in the olfactory bulb (47). Infection of these cells may help explain the loss of smell associated with some COVID-19 cases. However, the virus may also gain access via disruption of the blood brain barrier, as these were inflamed in most animals and neurovasculitis has been found in humans (48). Overall, the K18-hACE2 system may help shed light on this poorly studied area of human disease.

In addition to the application of K18-hACE2 model for the evaluation of MCMs, we conclude that this model may also be important in the study of SARS-CoV-2 pathogenesis. The identification of host factors driving the severe pathogenic processes in the lung, such as the role of macrophages and IL-6 in driving lung damage may provide critical insight into the molecular and cellular determinates of COVID-19 pathogenesis. Furthermore, as neurological sequelae during COVID-19 are an important component to severe disease, the K18-hACE2 model may provide a more defined understanding of SARS-CoV-2 interaction with the CNS.

## MATERIALS AND METHODS

### Viruses and cells

A third passage SARS-CoV-2 strain WA-1/2020 viral stock was obtained from the CDC and is from a human non-fatal case isolated in January 2020. A master challenge stock of virus was propagated by making two passages in Vero76 cells in Modified Eagles Medium with Earle’s Salts (EMEM) (Corning) supplemented with 1% GlutaMAX, 1% NEAA, and 10% heat inactivated fetal bovine serum (FBS) (Gibco). After 3 days, supernatant was collected and clarified by low speed centrifugation. Virus (P5 from the founder stock of virus) was quantified by plaque assay and determined to be endotoxin free. All virus work was handled in BSL-3 containment at USAMRIID.

### Mice

C57BL/6J (BL6), Rag2 KO mice and K18-hACE2 mice [B6.Cg-Tg(K18-hACE2)2Prlmn/J] (6-8 weeks old) were purchased from the Jackson Laboratory. Mice under isoflurane anesthesia were challenged intranasally with the indicated dose of SARS-CoV-2 WA-1/2020 strain diluted in a total volume of 50 μL of PBS (25 μL per nare). Mice were uniquely identified using ear tags (Stoelting) and provided an acclimation period of at least one week prior to commencement of the experiment. Mice were supplied nutrient gel when weights began to decrease.

### Ethics statement

All animal studies were conducted in compliance with the Animal Welfare Act and other federal statutes and regulations relating to animals and experiments involving animals and adheres to principles state in the Guide for the Care and Use of Laboratory Animals, National Research Council(49). Animal experimental protocols were approved by a standing internal institutional animal care and use committee (IACUC). The facilities where this research was conducted are fully accredited by the Association for Assessment and Accreditation of Laboratory Animal Care International. Animals meeting pre-established criteria were humanly euthanized after consultation with veterinary staff. Despite efforts to euthanize moribund mice, some were found dead (n=2).

### RT-qPCR

Organ tissue was homogenized in 750 μl Trizol reagent using a Tissuelyser II (spleen and liver) (Qiagen, Germantown, MD) or gentleMACS dissociator system on the RNA setting (Lung). RNA was extracted from Trizol per with modifications to the manufacturer’s protocol. Briefly, following the addition of chloroform and transfer of the aqueous layer, equal amounts of 70% Ethanol (Decon Laboratories, Inc) was added to the aqueous layer, then the nucleic acid was purified using the RNeasy Mini Kit (Qiagen) according to the manufacturer’s instructions. A Nanodrop 8000 was used to determine RNA concentration, which was then raised to 100ng/μl in UltraPure distilled water. Samples were run in duplicate, 500 ng per sample, on a BioRad CFX thermal cycler using TaqPath 1-step RT-qPCR master mix according to the CDC’s recommended protocol of 25°C for 2 minutes, 50°C for 15 minutes, 95°C for 2 minutes, followed by 45 cycles of, 95°C for 3 seconds and 55°C for 30 seconds. The forward and reverse primer and probe sequences are: 2019-nCoV_N2-F, 5’-TTA CAA ACA TTG GCC GCA AA-3’, 2019-nCoV_N2-R, 5’-GCG CGA CAT TCC GAA GAA-3’, and 2019-nCoV_N2-P, 5’-ACA ATT TCC CCC AGC GCT TCA G-3’. The limit of detection for this assay is 1 ×10^4^ copies. A synthetic RNA containing the real-time RT-PCR assay target sequence was acquired from Biosynthesis, Inc. (Portland, OR,USA). The approximate target copy number was determined using the nucleic acid sequence and the molecular weight of the synthetic RNA, and a stock at a known target copy number per ml was generated by resuspending the RNA in TE buffer. A standard curve was generated by serially diluting the stock standard curve in DNAse/RNAse free water (Invitrogen) and running the curve concurrently with the samples being tested.

### Histology

Necropsy was performed on the indicated organs. Tissues were immersed in 10% neutral buffered formalin for 14 days. Tissue were then trimmed and processed according to standard protocols (50). Histology sections were cut at 5-6 μM on a rotary microtome, mounted onto glass slides and stained with hematoxylin and eosin (H&E). Examination of the tissue was performed by a board-certified veterinary pathologist.

### Cytokine and chemokine analysis

Serum cytokine and chemokine analysis was performed using a magnetic bead-based plex mouse panel (Thermo fisher) targeting the indicated molecules. 25 μl of serum per mouse per time point was used. Plates were analyzed on a MAGPIX system (Millipore Sigma) and quantitated against standard curves using MILLIPLEX analyst software.

### *In situ* hybridization

To detect SARS-CoV-2 genomic RNA in FFPE tissues, in situ hybridization (ISH) was performed using the RNAscope 2.5 HD RED kit (Advanced Cell Diagnostics, Newark, CA, USA) as described previously(51). Briefly, forty ZZ ISH probes targeting SARS-CoV-2 genomic RNA fragment 21571-25392 (GenBank #LC528233.1) were designed and synthesized by Advanced Cell Diagnostics (#854841). Tissue sections were deparaffinized with xylene, underwent a series of ethanol washes and peroxidase blocking, and were then heated in kit-provided antigen retrieval buffer and digested by kit-provided proteinase. Sections were exposed to ISH target probe pairs and incubated at 40°C in a hybridization oven for 2 h. After rinsing, ISH signal was amplified using kit-provided Pre-amplifier and Amplifier conjugated to alkaline phosphatase and incubated with a Fast Red substrate solution for 10 min at room temperature. Sections were then stained with hematoxylin, air-dried, and coverslipped.

### Immunofluorescence

Formalin-fixed paraffin embedded (FFPE) tissue sections were deparaffinized using xylene and a series of ethanol washes. After 0.1% Sudan black B (Sigma) treatment to eliminate the autofluorescence background, the sections were heated in Tris-EDTA buffer (10mM Tris Base, 1mM EDTA Solution, 0.05% Tween 20, pH 9.0) for 15 minutes to reverse formaldehyde crosslinks. After rinses with PBS (pH 7.4), the section were blocked with PBT (PBS +0.1% Tween-20) containing 5% normal goat serum overnight at 4°C. Then the sections were incubated with primary antibodies: rabbit polyclonal anti-SARS-CoV Spike at a dilution of 1:200 (40150-T62-COV2, Sino Biological, Chesterbrook, PA, USA), mouse monoclonal anti-SARS-CoV NP at a dilution of 1:200 (40143-MM05, Sino Biological), mouse monoclonal anti-pan cytokeratin at a dilution of 1:100 (sc-8018, Santa Cruz Biotechnology, Dallas, TX, USA), mouse monoclonal anti-e-cadherin at a dilution of 1:100 (33-4000, Thermo Fisher Scientific, Waltham, MA, USA), rabbit polyclonal anti-<u>m</u>yeloperoxidase (MPO) at a dilution of 1:200 (A039829-2, Dako Agilent Pathology Solutions, Carpinteria, CA, USA), rabbit polyclonal anti-CD3 antibody at a dilution of 1:200 (A045229-2, Dako Agilent Pathology Solutions), rat monoclonal anti-CD45 antibody at a dilution of 1:100 (05-1416, Millipore Sigma, Burlington, MA, USA), rabbit polyclonal anti-CD68 at a dilution of 1:200 (ab125212, Abcam, Cambridge, MA, USA), mouse monoclonal anti-NeuN at a dilution of1:200 (MAB377, Millipore Sigma), and/or chicken polyclonal anti-GFAP at a dilution of 1:200 (ab4674, Abcam) for 2 hours at room temperature. After rinses with PBT, the sections were incubated with secondary goat anti-rabbit or anti-chicken Alexa Fluor 488 at dilution of 1:500 (Thermo Fisher Scientific) and goat anti-mouse or anti-rat Alexa Fluor 568 at a dilution of 1:500 (Thermo Fisher Scientific) antibodies, for 1 hour at room temperature. Sections were cover slipped using the Vectashield mounting medium with DAPI (Vector Laboratories). Images were captured on a Zeiss LSM 880 confocal system (Zeiss, Oberkochen, Germany) and processed using ImageJ software (National Institutes of Health, Bethesda, MD).

### TUNEL staining

Formalin-fixed paraffin embedded (FFPE) lung tissue sections were deparaffinized in Xyless II buffer (LabChem) 2x for 10 m each and rehydrated using an ethanol gradient (100%, 95% and 70%). Apoptotic cells were detected using the DeadEnd Fluorometric TUNEL assay (Promega) according to manufacturer’s protocol.

### NanoString gene expression analysis

Total RNA samples were analyzed using the nCounter mouse neuroinflammation (v1.0), nCounter mouse neuropathology (v1.0) and nCounter mouse myeloid (v2.0) code sets. Probe set-target RNA hybridization reactions were performed according to the manufacturer’s protocol (NanoString). For each hybridization reaction, 100 ng total RNA was used. Purified probe set-target RNA complexes from each reaction were processed and immobilized on nCounter cartridges using an nCounter Max preparation station, and transcripts were quantified on a digital analyzer (Gen, v.2). Data from each NanoString panel were first processed independently using nSolver (v.4.0) software (NanoString) as follows: following quality control checks on the individual RCC files, raw counts across samples were normalized to the mean counts for spiked synthetic DNA-positive controls present in the hybridization reactions to mitigate platform-associated sources of variation. Background thresholding was performed to the mean of negative control samples. Candidate reference genes were selected using the nCounter advanced analysis (nCAA) module (v.2.0.115), which implements the geNorm algorithm for downselection(52). Starting with a set of candidate reference genes, the algorithm identified the top five most stable genes for each panel. For each sample, normalization was performed by dividing the counts for each gene by the geometric mean for the five selected reference genes. These three normalized data sets were then combined in nSolver as a multi-RLF merge experiment and then input into the nCAA module for differential expression, gene set, cell type and biological pathway analysis. The threshold for differential expression was a log2 fold change of >1 and a p-value of <0.05.

### Statistical analysis

Survival statistics utilized the log-rank test. Statistical significance of virus titers were determined using the unpaired two-tailed Student’s *t* test. Significance levels were set at a *p* value less than 0.05. All analyzes were performed using Prism software.

## ACKNOWLEDGEMENTS

This project was funded by a grant awarded to J.W.G. and J.W.H. from the Military infectious disease program. We thank the USAMRIID histology lab and comparative medicine division for their assistance. We also express gratitude to the Jackson Laboratory for providing early access to the K18-hACE2 mice. Opinions, interpretations, conclusions, and recommendations are ours and are not necessarily endorsed by the U.S. Army or the Department of Defense.

## Author Contributions

J.W.G. and J.W.H. designed the studies. J.W.G, C.R.C, X.Z., A.R.G., B.D.C., E.M.M., L.E.W., J.D.S., R.L.B., J.L., A.M.B., H.B.R., J.M.S., B.S.H., C.F., and C.V.B performed research. J.W.G, C.R.C. and X.Z. analyzed data. J.W.G., C.R.C. and J.W.H. wrote initial drafts of the manuscript. All authors contributed to the final manuscript.

## Supplementary Materials

### Supplemental Materials and Methods

#### Duplex *In situ* hybridization

Duplex in situ hybridization was performed using the RNAscope 2.5 HD Duplex Assay kit (Advanced Cell Diagnostics) according to the manufacturer’s instructions with minor modifications. In addition to SARS-CoV-2 genomic RNA probe mentioned above (#854841, green), another probe with C2 channel (#848031-C2, red) specifically targeting human ACE2 (NM_021804.3) was designed and synthesized by Advanced Cell Diagnostics. ISH signal was amplified using kit-provided Pre-amplifiers and Amplifiers conjugated to either alkaline phosphatase or horseradish peroxidase, and incubated sequentially with a Fast Red and green chromogenic substrate solution for 10 min at room temperature. Sections were then stained with hematoxylin, air-dried, and coverslipped.

**Figure S1.**
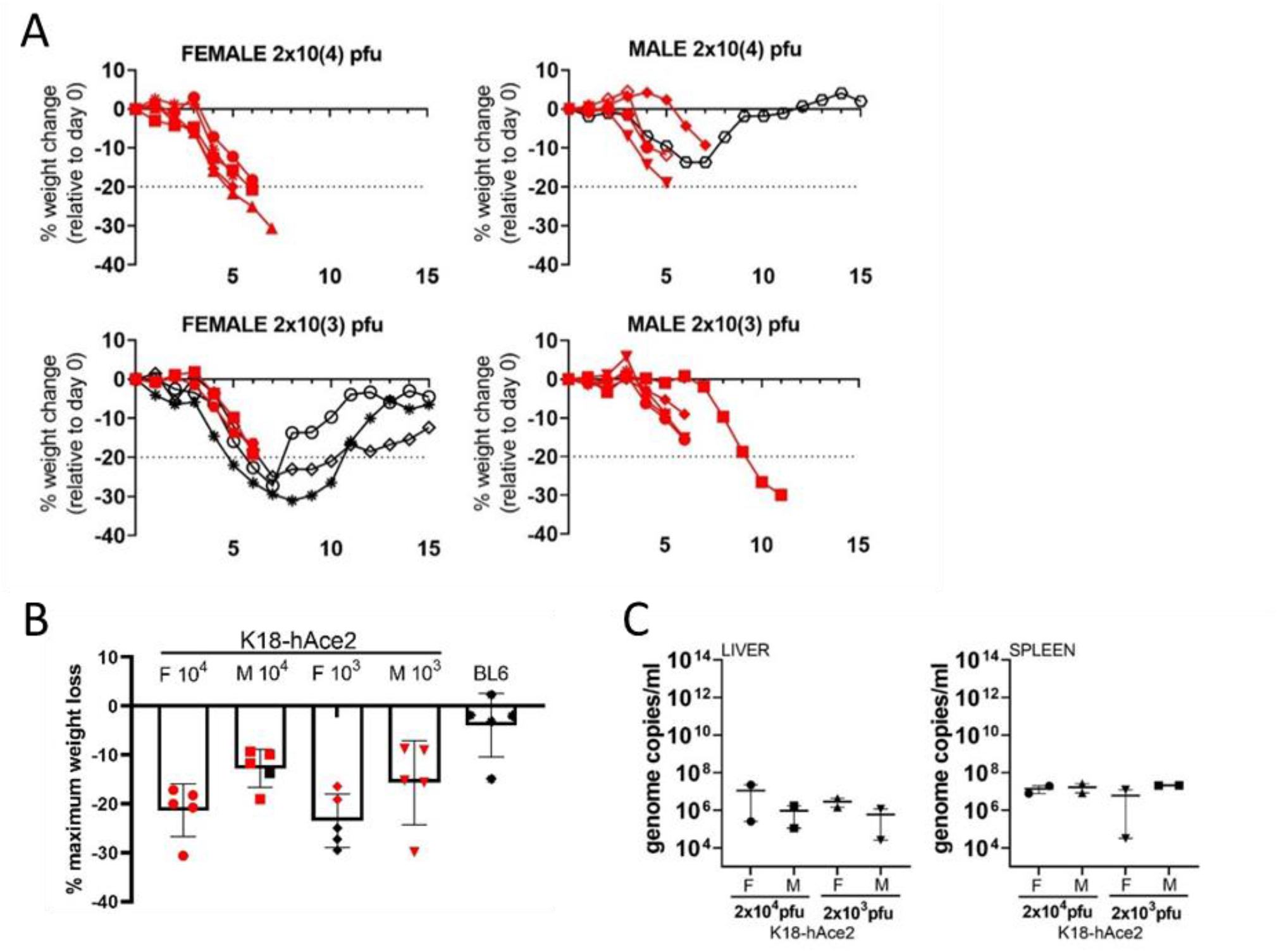
SARS-CoV-2 infection in K18-hAce2 transgenic mice. **A**. Individual weight loss in each challenge group is shown up to day 15. Red indicates an animal that died or was euthanized. **B**. Maximal weight loss over 15 day in infected mice. Red indicates mice that succumbed to disease. **C**. Titers in liver and spleen (n=2 mice/group) were examined on day 3 by qRT-PCR. Mean titers +/− SEM of the genome copies/ml were graphed.

**Table S1:** Summary of microscopic histopathology findings.

| **Organ** | **Microscopic Finding** | **n** | **Severity\*** |
| --- | --- | --- | --- |
| Lung | Vasculitis in small to intermediate size vessels | 21/22 | minimal-moderate |
|  | Alveolar septal inflammation / thickening | 20/22 | minimal-moderate |
|  | Perivascular inflammation (in vessels without vasculitis) | 15/22 | minimal-moderate |
|  | Alveolar accumulation of mononuclear leukocytes | 10/22 | minimal-mild |
|  | Alveolar exudate (fibrin, edema) | 9/22 | minimal-moderate |
|  | Type II pneumocyte hyperplasia | 8/22 | minimal-mild |
|  | Multinucleate cells (macrophages or viral syncytia) | 1/22 | minimal |
|  | Fibrin thrombi | 1/22 | moderate |
|  | Positive In situ hybridization (ISH) for SARS-CoV-2 | 19/22 | mild-severe |
| Nasal Turbinates | Degeneration, atrophy or erosion of olfactory epithelium | 9/22 | minimal-mild |
|  | Exudate within nasal meatus | 2/22 | minimal |
|  | Positive In situ hybridization (ISH) for SARS-CoV-2 | 19/22 | minimal |
| Brain | Vasculitis (and/or perivascular inflammation) in small to intermediate sized vessels | 14/22 | minimal-moderate |
|  | Microgliosis surrounding/adjacent to vessels | 12/22 | minimal-moderate |
|  | Meningitis | 8/22 | minimal-moderate |
|  | Necrosis | 5/22 | minimal-mild |
|  | Fibrin thrombi | 2/22 | minimal |
|  | Perivascular hemorrhage | 1/22 | mild |
|  | Microgliosis, gray matter, multifocal | 1/22 | moderate |
|  | Positive In situ hybridization (ISH) for SARS-CoV-2 (note 11/15 positive in the olfactory bulb) | 15/22 | minimal-marked |

\*Severity scores for ISH and histologic findings are based on the following: Minimal = If 10% or less of the cells in the section are ISH-positive or are affected respectively; Mild = If between 11% and 25% of the cells in the section are ISH-positive or are affected respectively; Moderate = If between 26% and 50% of the cells in the section are ISH-positive or are affected respectively; Marked = If between 51% and 79 of the cells in the section are ISH-positive or are affected respectively; Severe = If between 80% or more the cells in the section are ISH-positive or are affected respectively.

**Figure S2.**
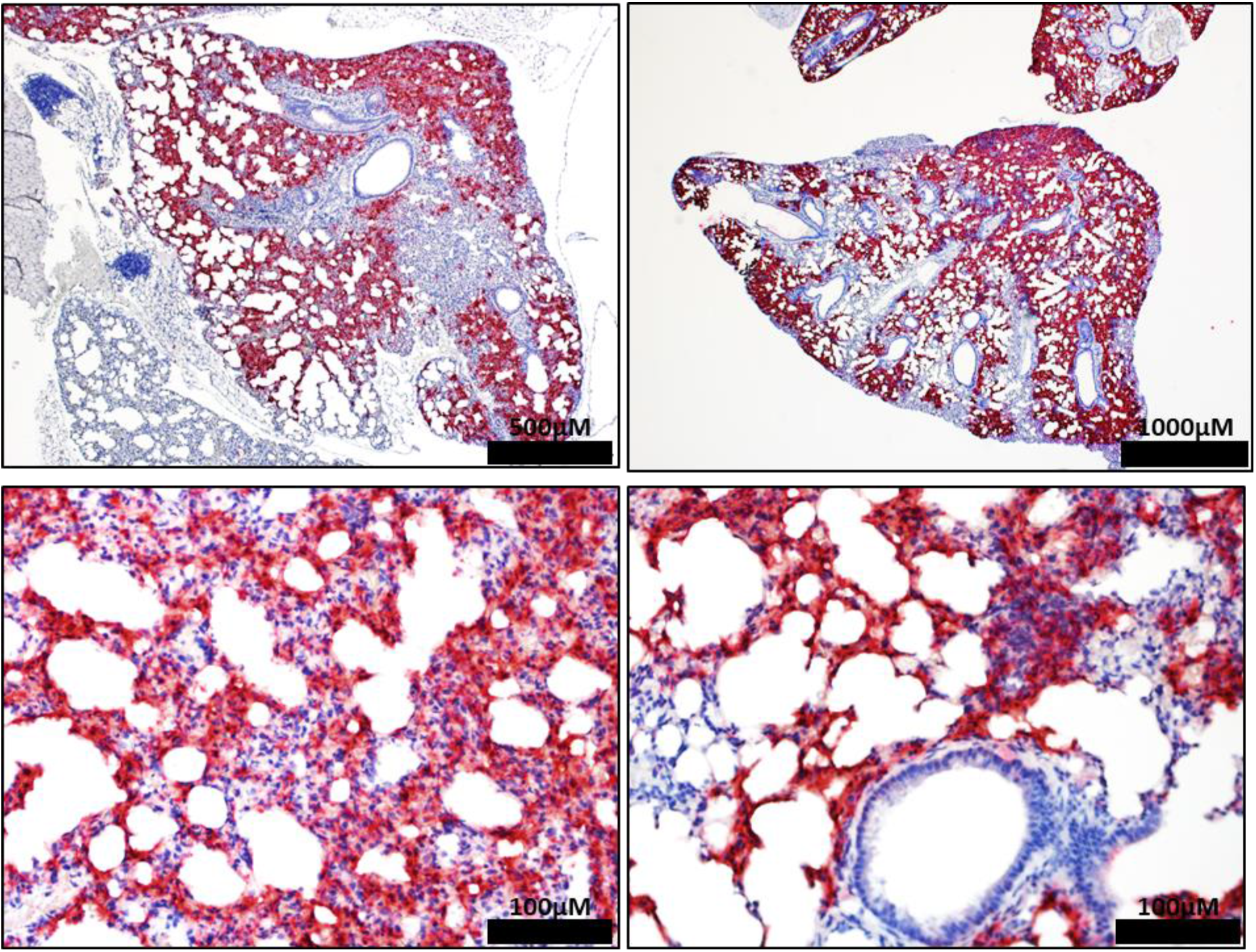
Infection of SARS-CoV-2 in the lungs in K18-Ace2 transgenic mice. Representative ISH images showing the presence of SARS-CoV-2 RNA (red) in the lungs of infected K18-hACE2 mice. ISH was performed in a different mouse than that in Fig. 2. Cells were counterstained with hematoxylin.

**Figure S3.**
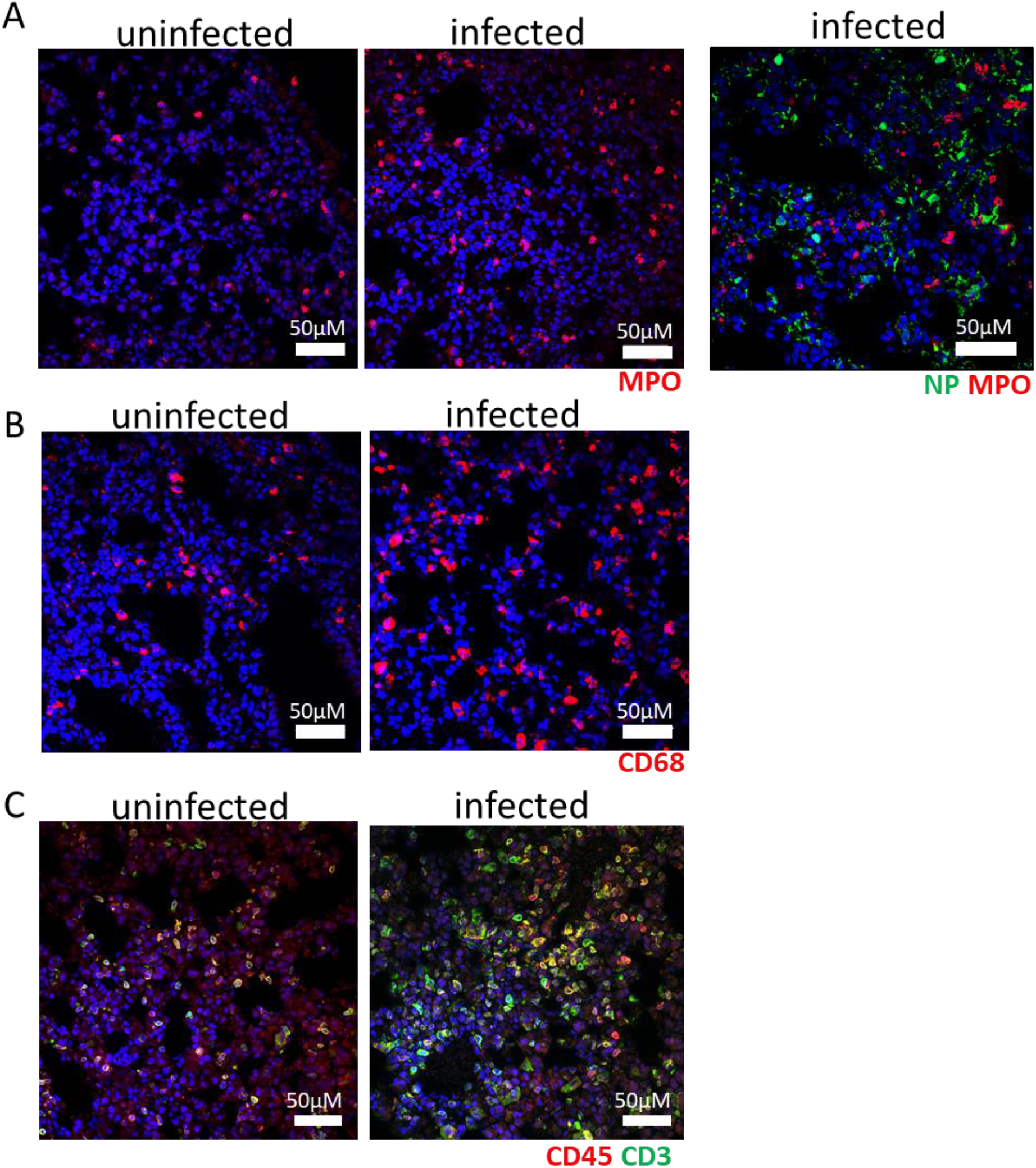
Infiltrating cells in the lungs of SARS-CoV-2 infected mice. **A–C.** IFA demonstrates increased number of myeloperoxidase (MPO)+ polymorphonuclear cells (neutrophils, eosinophils, and basophils) (A, red), CD68+ macrophages (B, red) CD45+ leukocytes (C, red) including CD3+ T cells (C, green) infiltrates in the lung of infected mice in comparison with the lung of uninfected mice. MPO positive cells (red) were devoid of viral NP protein (A, green). Nuclei are stained with DAPI (blue).

**Figure S4.**
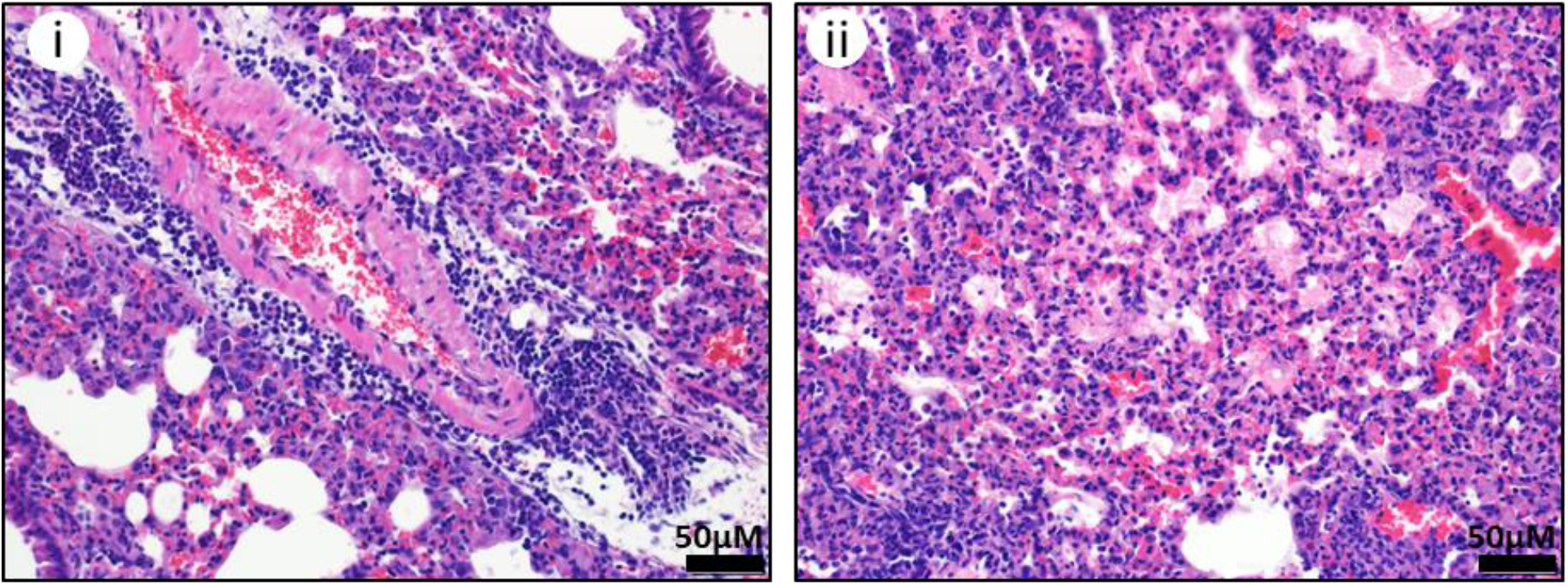
SARS-CoV-2 infection causes respiratory damage in K18-hACE2 mice. Representative H&E staining of lungs in infected K18-ACE2 mice. Edema, moderate numbers of mononuclear inflammatory cells, and fewer neutrophils expand the perivascular space surrounding an intermediate sized artery in the lung (i). Area of lung consolidation with inflammation/expansion of alveolar septa by fibrin, edema and mononuclear inflammatory cells; adjacent alveolar lumina are correspondingly filled with fibrin, edema and increased numbers of alveolar macrophages (ii).

**Figure S5.**
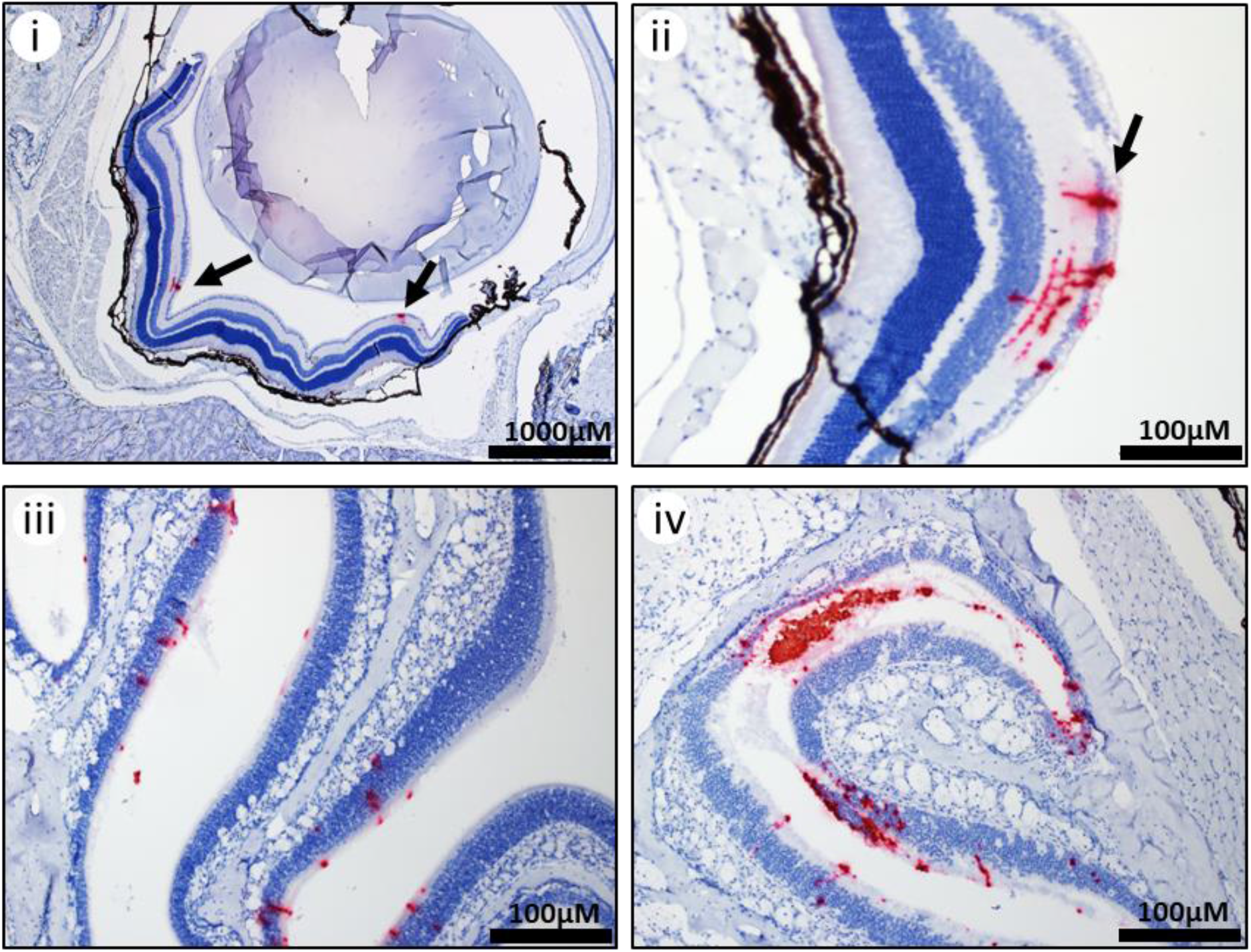
SARS-CoV-2 infection of eyes and nasal turbinates. Representative ISH staining showing the presence of SARS-CoV-2 RNA (red) in the eyes of infected K18-hACE2 mice with staining in the retina (arrows) (i & ii). Viral RNA was detected in the nasal turbiantes (iii & iv) with sloughing of infected cells (iv). Cells were counterstained with hematoxylin (blue).

**Figure S6.**
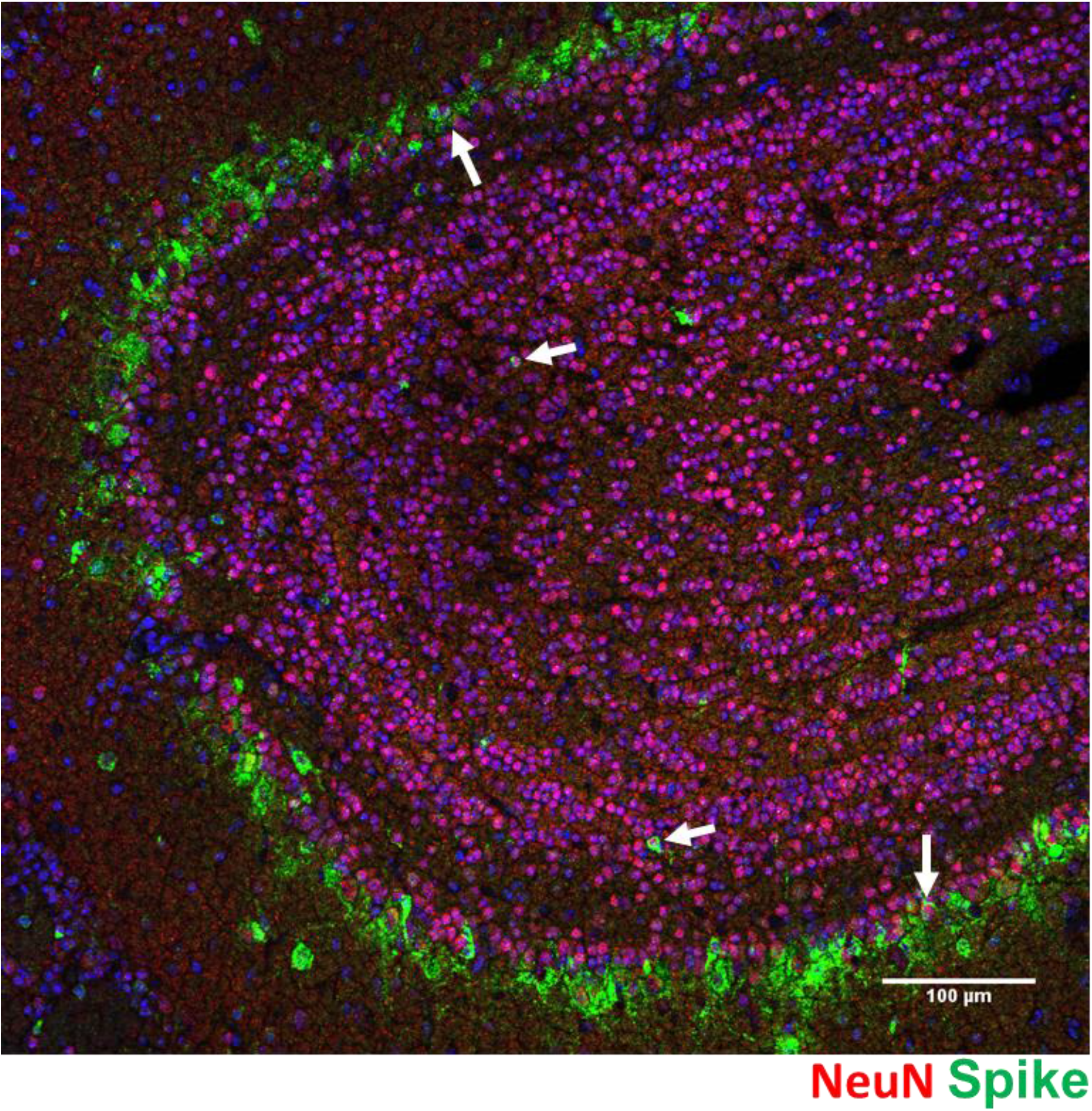
Infection of SARS-CoV-2 in the neurons of the olfactory bulb. Detection of viral spike protein (green) and the neuron marker NeuN (red) in infected olfactory bulb. Arrows denote co-stained cells. Nuclei are stained with DAPI (blue).

**Figure S7.**
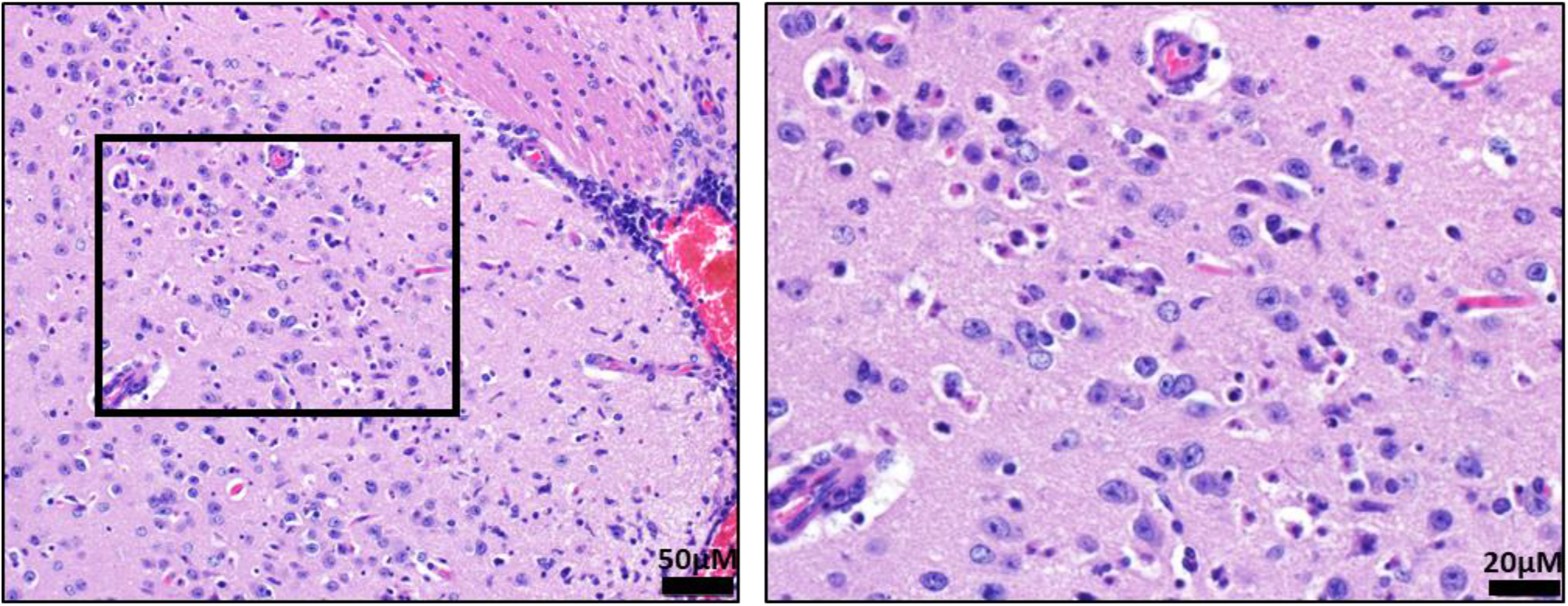
Brain lesions in SARS-CoV-2 infected mice. Representative H&E staining of lungs in infected K18-hACE2 mice. Multifocal areas of gliosis within the amygdala, predominantly characterized by increased numbers of microglia, and there are individual shrunken, angular cells with hypereosinophilic cytoplasm, pyknotic nuclei and surrounded by a clear halo, consistent with necrosis. While the morphology and location of individual necrotic cells is suggestive of neuronal necrosis, additional diagnostics are necessary to confirm the cell of origin. The right panel is a enhanced magnification of the boxed area

**Figure S8.**
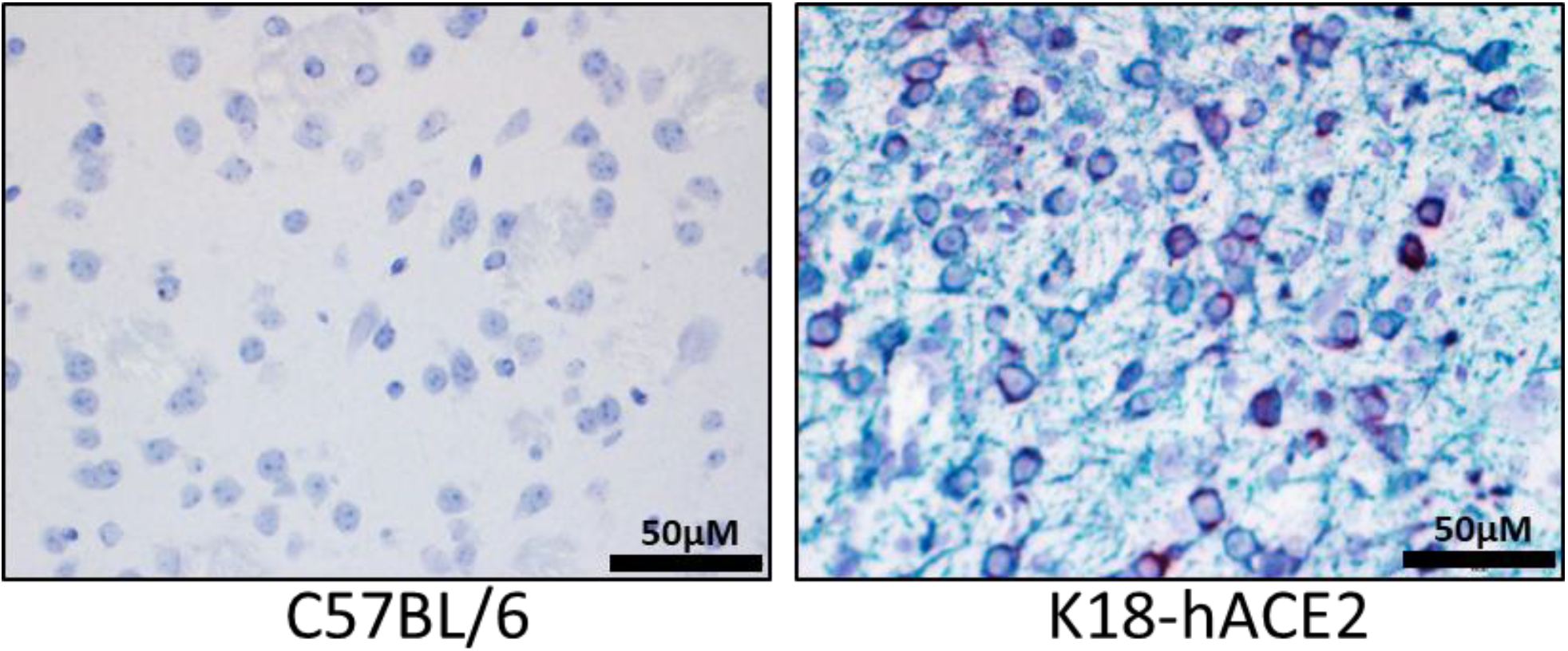
hACE2 transgene expression in neurons of SARS-CoV-2 infected mice. Representative ISH staining showing the presence of the hACE2 transgene (red) in infected K18-hACE2 mice. Duplex ISH staining further shows hACE2-exressping neurons are also positive to SARS-CoV-2 genomic RNA (green). Cells were counterstained with hematoxylin. C57BL/6 mice do not express the transgene.

